# Aphids capture plant inter- and intraspecific chemodiversity

**DOI:** 10.64898/2026.05.20.725638

**Authors:** Elizabeth Aiken, Selina Gaar, Jacqueline C. Bede, Caroline Müller, Thomas Dussarrat

## Abstract

The role of chemodiversity in plant-insect interactions is widely recognised. However, our understanding of the extent to which chemodiversity connects both partners remains limited. Here, we investigated how aphid chemistry is linked to their plant diet and whether aphids capture plant inter- and intraspecific chemodiversity. Up to 93% of aphid chemical features were detected in plants. Untargeted metabolomics of aphids feeding on diets composed of distinct species or chemotypes within species unveiled the aphid capacity to capture inter- and intraspecific chemodiversity. Multiple chemodiversity indices and metabolic features significantly tracked diet variation and plant chemotypes were reflected in aphid metabolites. These features included phenolics and amino acids, likely ingested with the phloem sap, and fatty acids and terpenoids, potentially captured from the leaf surface. Overall, these findings expand our knowledge of the aphid plant-derived chemical repertoire and highlight that plant chemodiversity can be transmitted, supporting the need for chemodiversity preservation programs.

## 1. Introduction

Co-evolution between plants and insects is governed by chemical principles that, *in fine*, influence ecosystem structure and dynamics (Dyer *et al*. 2018; Fraenkel 1959). The comprehension of these chemical principles is key to predicting the responses of trophic networks to climate change, preserving biodiversity, and supporting sustainable agriculture (Jamieson *et al*. 2017; Schuman *et al*. 2016). Extensive efforts have been conducted to explore and understand the consequences of plant-insect interactions on metabolites of both interacting partners. For example, previous work shed light on plant responses to herbivory (Schuman & Baldwin 2016) and other studies highlighted fascinating examples of how some insect species exploit plant chemical resources (Opitz & Müller 2009; Wheat *et al*. 2007). Conversely, a holistic understanding of how chemical diversity itself, beyond individual compounds, connects across kingdoms is still missing (Meiners 2015). Capturing these connections, as previously done at the biodiversity level (Karp *et al*. 2013; Scherber *et al*. 2010), is essential to uncover the hidden chemical networks that underpin ecosystem dynamics.

Plants have evolved a fascinating chemical diversity (hereinafter referred to as chemodiversity), which plays a central role in plant-environment interactions through the various functions of specific compounds or chemical blends (Dyer *et al*. 2018; Hanusch *et al*. 2026). Recently, plant chemodiversity, which refers to the richness, diversity and disparity of metabolites, was acknowledged as an important and dynamic niche dimension (Müller & Junker 2022). Interspecific chemodiversity has been characterised with the development of family-specific defensive chemicals linked to insect community composition (Agrawal *et al*. 2012; Richards *et al*. 2015). Although described decades ago (Stahl & Jork 1964), intraspecific chemodiversity has only recently gained prominence, and its significance in governing insect preferences and performance has been demonstrated. For example, glycoalkaloid composition of *Solanum dulcamara* was linked to slug resistance (Calf *et al*. 2018), while variation in the terpenoid profile of *Tanacetum vulgare* (synonym *Chrysanthemum vulgare*) influenced aphid preference (Jakobs & Müller 2018). However, most studies have focused on the consequences of individual chemotypes (*i*.*e*. groups of plants within a species that express a distinct profile of metabolites belonging to one chemical family (Müller *et al*. 2026)), whereas insects are exposed to a diverse range of chemotypes in natural environments (Kleine & Müller 2011). While a few studies investigated the impact of intraspecific chemodiversity on insect communities at the plot scale (Ojeda-Prieto *et al*. 2025; Ziaja & Müller 2023), its functioning at the ecosystem scale remains largely unknown (Hanusch *et al*. 2026).

The influence of plant diet on insect defence and reproduction has been previously described with the focus on specific metabolites. For instance, the plant diet influenced the insect cuticular hydrocarbon profile, which was linked to survival (Lima *et al*. 2024). Monarch butterflies captured cardenolides when consuming tropical milkweed for their own benefit, and Aulacoscelinae beetles exude host plant cycad-derived azoxyglycosides in reflexive bleeding from the leg joints when threatened (Agrawal *et al*. 2021; Prado *et al*. 2011). Phloem sap feeders also exploit plant chemicals. For example, milkweed bugs and the aphid species *Brevicoryne brassicae* sequester cardiac glycosides and glucosinolates, respectively, as defence against predators (Kos *et al*. 2012; Petschenka *et al*. 2022). While these findings have effectively revealed the use and fate of specific chemicals, a more general understanding of the extent to which the insect metabolome is shaped by its diet remains to be elucidated. Addressing this challenge requires comprehensive analytical approaches, and untargeted metabolomics has emerged as an ideal tool in this context (Díaz *et al*. 2024).

Here, we assessed how the inter- and intraspecific chemodiversity of the plant diet influences aphid chemistry. More specifically, we combined multivariate statistics and untargeted metabolomics to (i) explore the chemical connectedness between plants and aphids from chemical families to the metabolite level and (ii) evaluate the aphid capacity to capture plant inter- and intraspecific chemodiversity. We hypothesised that variation in aphid chemodiversity reflects the dietary plant species composition. In addition, we hypothesised that chemodiversity principles, such as the existence of chemotypes, extend to the aphid metabolome. This hypothesis was supported by studies showing that phloem sap-sucking insects can take up diverse compounds (*e*.*g*. glucosinolates) and that some insects, such as ants, harbour colony-specific chemical signatures (Martin *et al*. 2013; Sun *et al*. 2021). To test our hypotheses, two experiments were conducted. The first explored the impact of a change in species composition of the plant diet on the feeding specialist *Macrosiphoniella tanacetaria* and the generalist aphid *Myzus persicae* after 24 hours and one or two generations. The second explored the consequences of a change in plant species or chemotype composition within a plant species on the metabolic composition of *M. tanacetaria* using the highly chemodiverse plant species *T. vulgare* and a close relative. Putative annotation of chemical families and metabolites significantly influenced by variation in the plant diet provided insights into whether aphids efficiently capture the diet’s chemodiversity. Finally, we briefly discuss the potential consequences of the findings at the ecosystem scale.

## 2. Materials and Methods

### 2.1. Plant and aphid material

Five *T. vulgare* chemotypes defined by their leaf terpenoid profile, “BThu”, dominated by β-thujone; “ABThu”, dominated by α- and β-thujone; “Keto”, dominated by artemisia ketone; “Aacet”, dominated by artemisyl acetate, artemisia ketone, and artemisia alcohol; and “Myrox”, dominated by (*Z*)-myroxide, santolina triene, and artemisyl acetate (Ziaja & Müller 2023), were used in this study. *T. vulgare* plants were grown from cuttings taken from our *T. vulgare* stock, which was established using seeds collected in 2019. Pots of *Chrysanthemum indicum*, a related species to *T. vulgare* (both Asteraceae) on which *M. tanacetaria* can also feed, were bought at the market. *Brassica oleracea* (cabbage) and *Pisum sativum* (pea) were grown from commercially purchased seeds. Plants were grown in the greenhouse and transferred to a climate chamber (16-hour light / 8-hour dark photoperiod with 23°C/18°C and 60-70% relative humidity) at least three weeks before the experiment, watered twice weekly. Plants of *T. vulgare* and *C. indicum* were used during flowering time, while *B. oleracea* and *P. sativum* were used before flowering.

Aphids of *M. tanacetaria* were collected from *T. vulgare* of the ABThu chemotype and subsequently reared on this chemotype in non-controlled laboratory conditions. A starting colony of the generalist aphid *M. persicae* was obtained from Göttingen University and reared on *B. oleracea*. Both species were kept for four weeks at room temperature in rearing tents (60cm*60cm*60cm, Bugdorm BD2F120) in the laboratory prior to the experiment. During this period, both aphid species reproduced extensively.

### 2.2. Design of the “Generation” and “Diversity” experiment and sampling

To explore the chemical links between plant diet and aphids, two experiments were conducted (Fig. S1). The first experiment, referred to as the “Generation” experiment, evaluated the extent to which aphid metabolic composition is affected by a change in species composition of the plant diet. Nymphs of *M. tanacetaria* and *M. persicae* were placed on potted plants of *B. oleracea* or ABThu (*T. vulgare* chemotype), respectively, at time T0 in tents in the climate chamber (one hundred nymphs per plant, four replicates per condition). At time T0, 30 aphids were sampled, 40-50 aphids were transferred from *B. olearacea* to *P. sativum* or from *T. vulgare* to *C. indicum* in separate tents or kept on their original diet, as a control. After 24 hours, approximately 30 aphids were sampled in both original and new diet conditions (T24h). Furthermore, 30 aphids of the offspring (“Progeny 1”, T2 weeks) were collected per condition two weeks after the initial transfer, and of the second generation after four weeks (“Progeny 2”, T4 weeks). At this last time point, plants showed signs of senescence. Consequently, only three or two replicates of *M. tanacetaria* feeding on *T. vulgare* or *C. indicum* for four weeks could be collected, respectively, as well as only one replicate for *M. persicae* feeding on *P. sativum* and zero on *B. olearacea*. In addition, four to five replicates of the third and fourth youngest leaf from each plant species used in this experiment (plus plants of Keto and Myrox chemotypes) were sampled from undamaged flowering plants kept under the same growing conditions (Table S1). For sampling, leaves and aphids were collected into 2 mL Eppendorf tubes (aphids with paintbrushes), directly snap-frozen in liquid nitrogen, and stored at −70°C.

The second experiment (“Diversity” experiment) aimed to explore the chemical consequences on aphids of a change in plant species or chemotype composition within the same plant species in the diet (Fig. S1B). Sixty nymphs of *M. tanacetaria* were placed into each of four feeding conditions, each comprising four plants on which the aphids fed sequentially: (i) C1, four plants of the same chemotype (eight replicates with Aacet chemotype and eight with BThu), (ii) C2, two plants of two chemotypes (Aacet and BThu), (iii) C3: one plant of four distinct chemotypes (Aacet, BThu, Keto, and Myrox), and (iv) C4: one plant of three distinct chemotypes (Aacet, BThu, and Keto) plus one plant of *C. indicum*. Aphids were fed for two days on plant 1 enclosed in a mesh bag (14 cm x 14 cm), and then transferred to the next plant. Between each transfer, the potential aphid progeny was removed, and the leaf on which aphids fed was sampled. After 8 days (4 plants x 2 days), aphids were sampled. Therefore, for each replicate, one aphid sample (composed of approximately 30 adult aphids) and its respective diet (one leaf of each of the plants on which they fed) were obtained. In total, 16 replicates for C1 and C2, 18 replicates for C3 and 14 replicates for C4 were obtained. However, one replicate of C4 exploded at sampling due to liquid nitrogen and could not be analysed (Table S1). Between replicates, the feeding order and the leaf age on which aphids fed (third to sixth leaf) were randomised (*e*.*g*. for C3, a Keto plant could be the first, second, third or fourth feeding source), and each aphid fed on plants of four distinct maternal origins. As plants of multiple chemotypes can originate from the same maternal plant (Dussarrat *et al*. 2023), the same maternal origin can be observed in different chemotypes (*e*.*g*. a Keto plant and a BThu plant can have the same maternal origin). Aphid and leaf sampling and storage were performed as described above. Samples were then freeze-dried for 48 hours, ground, and weighed for chemical extraction.

### 2.3. Chemical extraction

To 12 mg of plant samples, 400 µL of a 90% methanol solution (v:v) with hydrocortisone (10 mg/L) as internal standard (Sigma-Aldrich, Steinheim, Germany), while 250 µL was added to each of the entire aphid samples. As the number and weight of aphids could vary between samples, the dry weight per sample was noted before grinding for further normalisation. The mixture was vortexed for 5 min and sonicated in an ice bath for 20 min. After centrifugation, supernatants were collected and filtered using 0.2 µm syringe filters (Phenomex, Torrance, CA, USA), as previously described (Dussarrat *et al*. 2023).

### 2.4. Metabolomics analyses

Extracts were subjected to untargeted metabolic fingerprinting by UHPLC-QTOF-MS/MS (UHPLC: Dionex UltiMate 3000, Thermo Fisher Scientific, San José, CA, USA. QTOF: compact, Bruker Daltonics, Bremen, Germany) operating in negative mode via electrospray ionisation source. Separation was performed on a Kinetex XB-C18 column (150 × 2.1 mm, 1.7 µm, with guard column; Phenomenex) following the gradient detailed in Dussarrat *et al*. (2023), and a spectra rate of 5 Hz was used with optimised MS parameters. Extracts from both experiments were processed on LC-MS in the same sequence. The T-ReX 3D algorithm of MetaboScape (v. 2021b, Bruker Daltonics) was used for the preprocessing with the following settings: features in a minimum of 3 samples, intensity threshold 1000, and minimum peak length of 11 spectra. From the resulting table, peak intensities were normalised by the intensity of the internal standard and sample dry weight. Quality controls (QC, mixture of 12µL from each extract) and extraction blanks that have been measured in the same sequence were used for subsequent data cleaning (ion kept if: average QC intensity > 5 times average blank intensity (i.e. peak height) and coefficient of variation in QC < 30%), yielding 3,874 and 4,288 features for the “Generation” and “Diversity” experiments, respectively. LC-MS datasets were normalised using median normalisation, cube-root transformation and Pareto scaling on MetaboAnalyst v.6 (Pang *et al*. 2021) prior to statistical analysis as previously described (Díaz *et al*. 2024, Tables S2-S5).

### 2.5. Putative annotations

Putative chemical formula and families and putative compound names were defined using SIRIUS (v. 6.1.0) and CANOPUS (Dührkop *et al*. 2015, 2019) with default parameters (10 ppm and de novo plus bottom up molecular formula generation with a threshold *m*/*z* of 500) and using the Natural Product Classification (NPC) ontology. Chemical families, which include chemical pathways, superclasses, and classes, were defined if the confidence score was ≥ 0.8, as recommended (Hoffmann *et al*. 2022). Confidence in the annotation was defined following the Metabolomics Standards Initiative confidence levels (MSI), with MSI 2 obtained only if both approximate confidence score and Tanimoto score ≥ 0.8 (Sumner *et al*. 2014). The richness, Shannon diversity, and Functional Hill diversity (total and for each chemical family) were calculated using the *chemodiv* R package, as previously described (Authier *et al*. 2026; Petrén *et al*. 2023). Finally, chemical families including less than 0.5% of the total detected features (19 and 21 chemical features for the Generation and Diversity experiment, respectively) were removed from the analyses as their biological relevance may be limited (Authier *et al*. 2026). Non-normalised and normalised (median normalisation, cube-root transformation and Pareto scaling) chemodiversity indices datasets from both experiments are available in Tables S6-S9. All chemical datasets and metadata are available online (see Data availability section).

### 2.6. Statistical analyses

Aphid and plant chemodiversity, as well as the metabolome of *M. tanacetaria* and *M. persicae*, were compared using Student’s t-tests in MetaboAnalyst (v. 6, Tables S10 and S11). Volcano plots were created using *ggplot* on R (v. 4.5.1) (R Core Team 2022; Wickham 2016). To visualise interesting clusters of chemical features varying between aphids and plants and between aphid species, feature-based metabolic networks were developed using GNPS with a paired cos score of 0.7 and a minimum of matched fragment ions of 5 (Nothias *et al*. 2020; Wang *et al*. 2016).

To test whether aphids capture interspecific chemodiversity, we used the normalised datasets of the “Generation” experiment. Principal component analyses (PCA), as well as supervised analyses such as partial least squares discriminant analyses (PLS-DA), were generated using MetaboAnalyst (v. 6). Significant chemical features and indices (*P* < 0.05, ANOVA test) were illustrated in barplots using *ggplot*.

Using data from the “Diversity” experiment, we explored the chemical variation induced by a range of diverse plant diets that included one to four *T. vulgare* chemotypes or three chemotypes plus *C. indicum*. As a first step to assess whether aphids can capture plant chemodiversity, we explored how increasing the number of chemotypes or the number of related plant species in a diet influenced its chemical diversity (*i*.*e*. plant diet level). We tested whether chemodiversity indices or chemical features could be used to differentiate *T. vulgare* chemotypes using PCA and sparse PLS-DAs. Plant chemodiversity indices and chemical features that showed significant variation between either chemotypes or diversity treatments (*i*.*e*. treatment classes C1 to C4) were extracted using ANOVA (*P* < 0.05). Next, the *vegan* package was used to calculate Bray-Curtis dissimilarities between diets of the same chemotype (*e*.*g*. two BThu diets), different chemotypes, or distinct treatments (*e*.*g*. C1 versus C2) with the “bray” method (Oksanen *et al*. 2022). Subsequently, we assessed the extent to which aphid chemodiversity varies between diets of different chemotypes or plant diversity levels using the same statistical approach. Finally, we explored the link between aphid chemical features that responded significantly to a change in diet composition (chemotype or diversity level) and plant chemistry using Pearson’s correlations via the *Hmisc* package (Jr 2025). For all tests, significance levels are available with and without FDR correction. R scripts used for statistical analyses were deposited online (see Data Availability section).

## 3. Results

### 3.1. Substantial overlaps between plant and aphid metabolome

Plants harboured a considerably more diverse metabolome than both aphid species, as shown by higher richness and FH diversity of the total detected metabolome, as well as of phenolics and terpenoids, for instance (Fig. 1A, Table S10). In contrast, both aphid species tended to show a higher FH diversity of a few specific lipid families, such as glycerophospholipids. Moreover, plants included 78% of the metabolome detected in aphids, while only 49% of the chemical features detected in plants were also observed in aphids (Fig. 1B). These results were supported by the Diversity experiment, where 93% of *M. tanacetaria* chemical features were also detected in plants (Fig. S2). Between the two aphid species, differences and commonalities were observed (Fig. 1C). A total of 1,558 chemical features differed between the two species (*P* < 0.05, Table S11 and Fig. S3). For example, glucosinolates were identified as a key discriminant chemical family with significantly higher abundance in *M. persicae* compared to *M. tanacetaria*. Fatty acid and phenolic pathways also showed pronounced variation between species, with different associated chemical classes showing contrasting patterns of abundance across aphid species (Fig. 1C).

**Fig. 1.**
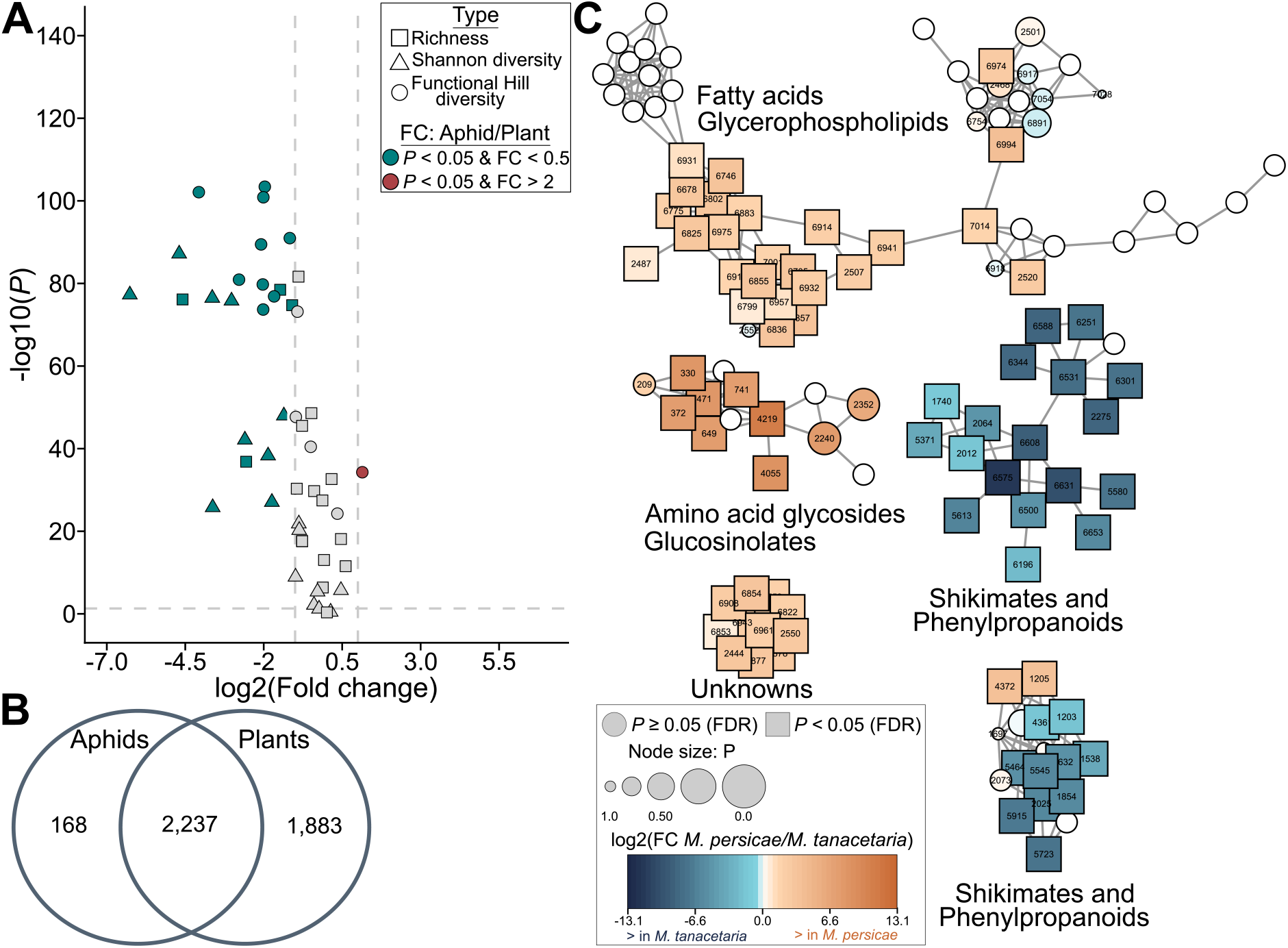
Comparison of aphid and plant chemistry in the “Generation” experiment. **A**. Depiction of richness, Shannon diversity and Functional Hill diversity of different chemical families in aphids (*Macrosiphoniella tanacetaria* and *Myzus persicae*) and leaves (*Brassica oleracea, Pisum sativum, Chrysanthemum indicum*, and *Tanacetum vulgare*). Fold changes (FC) were calculated as the ratio between aphids and plants. Blue represents higher values in plants (*P* < 0.05, fold change < 0.5) while red represents higher values in aphids. **B**. Numbers of detected metabolic features in aphids and plants. **C**. Cluster of compounds illustrating chemical variation between aphid species. Brown colour refers to higher contents in *M. persicae*, while blue colour refers to higher contents in *M. tanacetaria. FC*: fold change.

### 3.2. Aphids capture plant interspecific chemodiversity

To assess whether phloem sap-sucking insects can capture the interspecific chemodiversity from their diet, we compared the chemistry of *M. tanacetaria* feeding on the ABThu chemotype of *T. vulgare* or *C. indicum* and of *M. persicae* feeding on *B. oleracea* or *P. sativum*. An unsupervised analysis primarily highlighted the chemical variation between the two aphid species (Fig. 2A). Chemical consequences of the diet chemodiversity on the aphids were observable in chemodiversity indices, which could be used to partially distinguish aphids feeding on *T. vulgare* or *C. indicum* and to illustrate the plant species and generation effect on *M. persicae* (Fig. S4A-C). In fact, 32 and 20 chemodiversity indices responded to changes in dietary plant species composition in *M. tanacetaria* and *M. persicae*, respectively (Table S12). Notably, 60% and 36% of all chemical families analysed in this study responded to this diet, respectively (at least one chemodiversity index per family). Phenolic and amino acid-related pathways, as well as fatty acids, were influenced by the diet in both aphid species. In contrast, terpenoids were mostly influenced in *M. tanacetaria* (Fig. 2B).

**Fig. 2.**
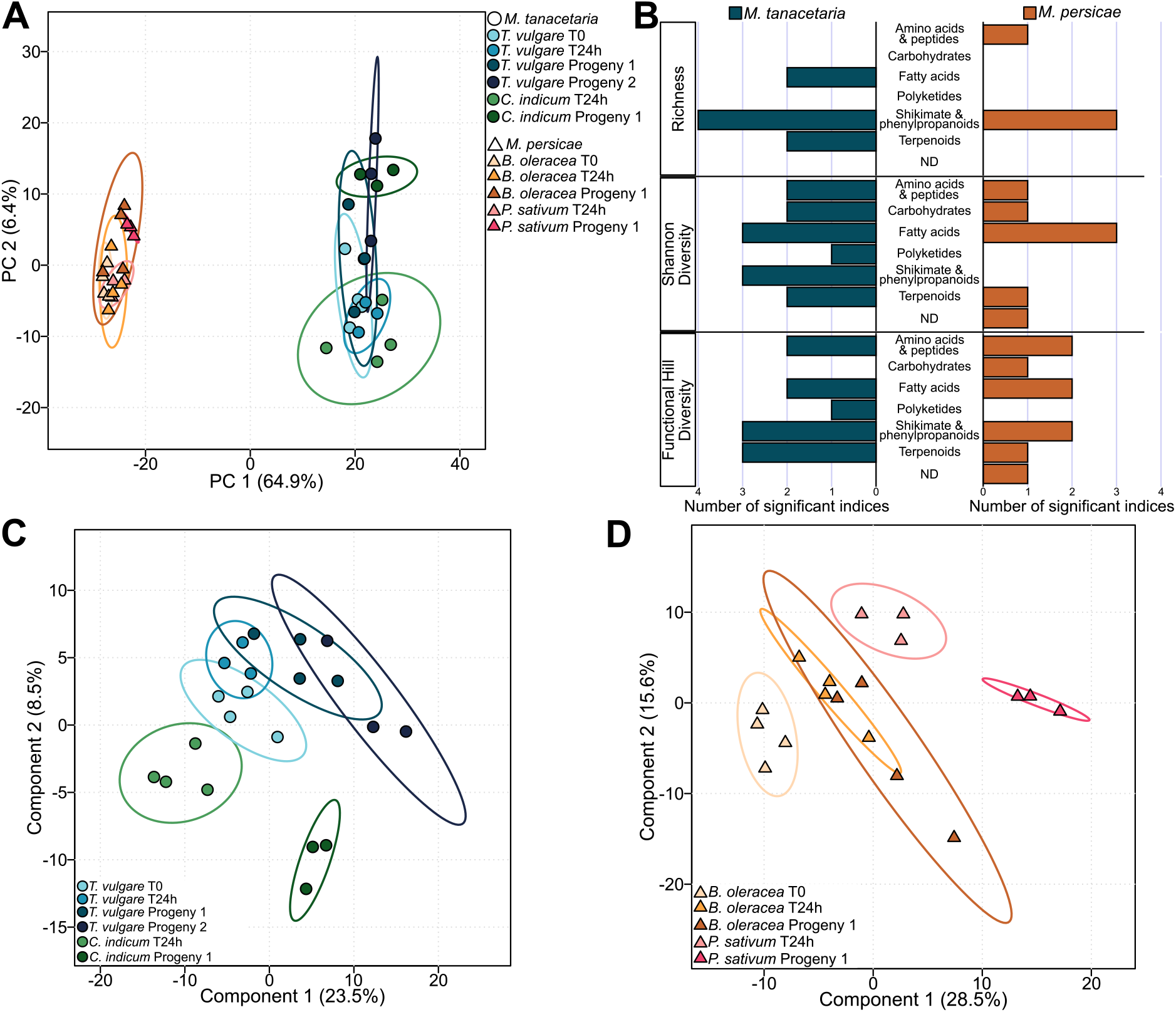
Aphids capture interspecific chemodiversity from the plant diet. **A**. Principal component analysis (PCA) of aphid metabolites from two species (*Macrosiphoniella tanacetaria* and *Myzus persicae*) feeding on distinct diets. **B**. Number of chemodiversity indices per chemical pathways influenced by the plant species included in the diet found in *M. tanacetaria* or *M. persicae* (*P* < 0.05). **C**. Partial least squares-discriminant analysis (PLS-DA) exploring the effect of diet on *M. tanacetaria* through asexual generations. R^2^ = 0.94, *P*_species_ = 0.063, *P*_generation_ = 0.001, *P*_species*generation_ = 0.085 via ANOVA test. **D**. PLS-DA exploring the effect of diet on *M. persicae* through asexual generations. R^2^ = 0.95, *P*_species_ = 0.002, *P*_generation_ < 0.001, *P*_species*generation_ = 0.070 via ANOVA test. *ND*: not determined.

At the metabolite level, PLS-DA analyses also highlighted that both aphid species effectively capture interspecific diversity (Fig. 2C and 2D). While changes at the metabolic level were observable after one day, the effect was getting more pronounced in the subsequent aphid generation(s). For instance, samples of *M. tanacetaria* feeding on *C. indicum* and *M. persicae* feeding on *P. sativum* showed a gradual shift away from their initial diets (*T. vulgare* T0 and *B. oleracea* T0, respectively) along the first component. A generation effect was also visible in aphids feeding on *T. vulgare* or *B. oleracea*. Overall, 606 aphid chemical features responded significantly to changes in dietary species composition. However, this chemical shift was aphid species-specific, with only 6% of significant features shared between both species (Fig. S4 and S5). Significant features included phenolics, carbohydrates, amino acids, and polyketides, and especially the occurrence of glucosinolates in *M. persicae* (Table S12).

### 3.3. Chemical redundancy in a diverse diet

To further characterise the capacity of aphids to capture the plant chemodiversity of their diet, we compared the chemistry of *M. tanacetaria* feeding on one to four *T. vulgare* chemotypes or three chemotypes plus a related species (*i*.*e. C. indicum*). Regarding the metabolic composition of the plant diets between diversity treatments, unsupervised and supervised analyses illustrated a clear distinction between the two plant species and, within *T. vulgare*, between the five chemotypes in the sPLS-DA analysis (Fig. 3A and S6). In contrast, PLS-DA failed to provide a clear distinction between diets composed of different *T. vulgare* chemotype combinations, but showed a significantly different pattern for the “C4” group, composed of three *T. vulgare* chemotypes plus *C. indicum* (Fig. 3B). This pattern was also reflected in Bray-Curtis distances. Chemical dissimilarities were higher when comparing diets from the distinct chemotypes or species than either the diets of the same chemotype or species or of distinct diversity treatments (Fig. 3C). This disparity was further supported by the number of chemical features that varied significantly between diets of different chemotypes or species and those from different diversity treatments, suggesting chemical redundancy when supplementing a diet with an additional chemotype or a related plant species (Fig. 3D, Table S13). In total, 85 and 40 plant chemodiversity indices and 2,370 and 1,090 features differed between chemotypes or species or between treatments, respectively (*P* < 0.05). For instance, diets of distinct chemotypes or species harboured distinct total richness and FH diversity, while both types of diet shift led to changes in the richness and FH diversity of terpenoid and fatty acid related pathways. At the metabolite level, chemical features influenced by the diet diversity covered all main chemical families, especially phenolics, terpenoids, and fatty acids (Table S13).

**Fig. 3.**
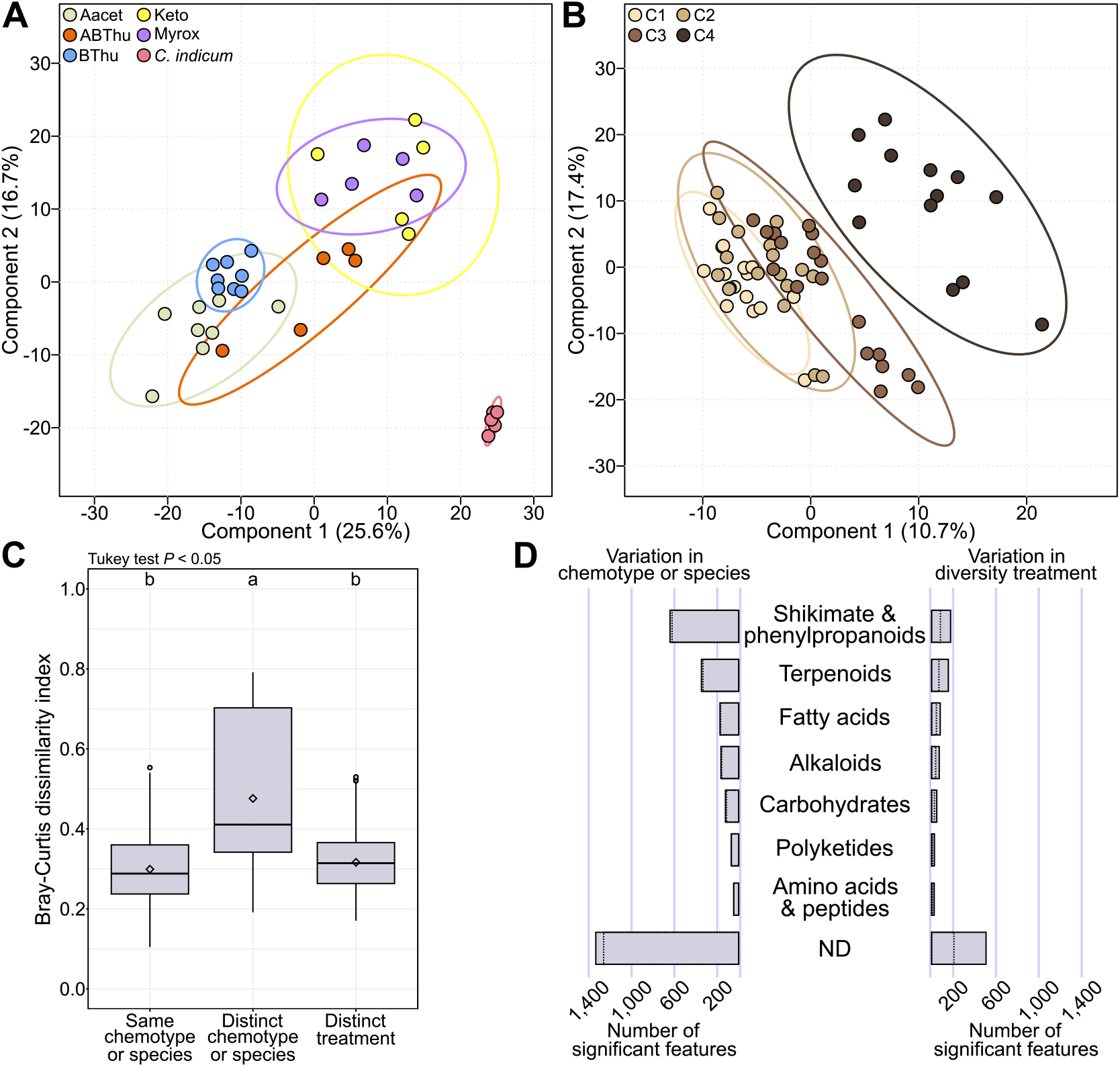
Chemical redundancy in a diverse diet. **A-B**. Partial least squares-discriminant analysis (PLS-DA) of leaf metabolites (4,288 features) comparing single chemotypes of *T. vulgare* or *C. indicum* (A, R^2^ = 0.73, *P* < 0.001 via ANOVA test) or treatments, *i*.*e*. diet mixes C1 to C4 (B, R^2^ = 0.78, *P* < 0.001 via ANOVA test). Diets included four plants of the same *Tanacetum vulgare* chemotype (C1), two plants of two chemotypes (C2), one plant of four distinct chemotypes (C3), and one plant of three distinct chemotypes plus one plant of the related species *Chrysanthemum indicum* (C4). **C**. Bray-Curtis dissimilarities between plant samples from “Same chemotype” (e.g. Keto-Keto or *C. indicum*-*C. indicum*), “Distinct chemotype or species” (Keto-Aacet, Keto-*C. indicum*) or “Distinct treatment” (e.g. C1-C2, C2-C3). **D**. Classification of features varying significantly between chemotypes or species or between treatments (*P* < 0.05) in plants. Dot bars represent significant features with *P* < 0.05 after FDR correction. *Aacet*: artemisyl acetate-artemisia ketone-artemisia alcohol chemotype, *ABThu*: α-/β-thujone chemotype, *BThu*: β-thujone chemotype, *Keto*: artemisia ketone chemotype, *Myrox*: (*Z*)-myroxide-santolina triene-artemisyl acetate chemotype. *ND*: not determined.

### 3.4. Aphids capture plant inter- and intraspecific chemodiversity

We subsequently examined whether chemical variation at the diet level was followed by corresponding changes at the aphid level (Fig. 4). Aphids efficiently captured plant intraspecific chemodiversity (Fig. 4 and S7). Plant chemotypes were reflected in aphid chemistry (Fig. 4A). Modifying the chemotypes present in the diet influenced a total of 18 chemodiversity indices from 10 chemical families and 229 chemical features, including phenolics, fatty acids and terpenoids, in the aphids (Fig. 4B and Table S13). For instance, Shannon and FH diversity of phenolics and terpenoids, as well as glycerophospholipid richness, varied significantly between aphids fed on different chemotype conditions. In addition, similar to observations at the plant level, varying diet diversity yielded a gradual shift from C1 (one chemotype) to C4 (three chemotypes plus *C. indicum*) in aphid features without a clear distinction between treatments when considering the overall composition of chemical features (Fig. 4C). However, 223 aphid chemical features responded to variation in diet diversity levels (Table S13). Pearson correlations between aphid chemistry and the chemistry of their respective diets indicated that these significant features were either positively or negatively correlated with the same feature in the diet (Fig. S7C). Significantly positively correlated features included diverse chemical families from primary and specialised metabolism, with a prevalence of phenolics (mostly flavonoids) and carbohydrates (Fig. 4D). Terpenoids (mostly monoterpenoids), amino acids, and fatty acids were also represented among the significant features (Fig. 4D and Table S14). Significantly negatively correlated features included similar chemical families, although polyketides were proportionally more represented.

**Fig. 4.**
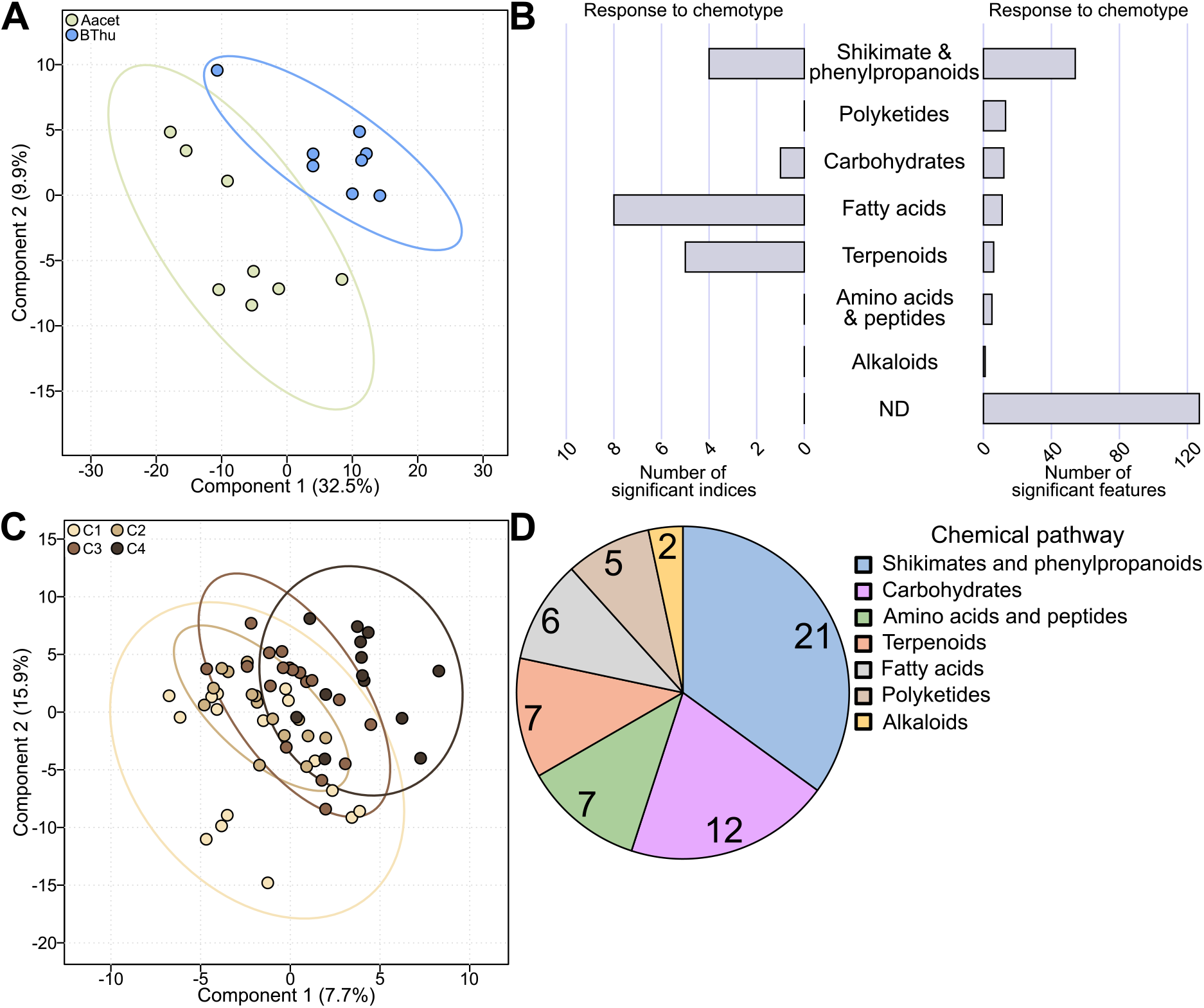
Aphids capture intraspecific chemodiversity from the plant diet. **A**. Partial least squares-discriminant analysis (PLS-DA) comparing aphid metabolism (4,288 features) that fed on the β-thujone chemotype (*BThu*) or the artemisyl acetate-artemisia ketone-artemisia alcohol chemotype (*Aacet)* of *Tanacetum vulgare*. R^2^ = 0.84, *P* = 0.048 (ANOVA test). **B**. Putative annotation of the chemodiversity indices and chemical features varying significantly in aphids between Aacet or BThu diet (*P* < 0.05). **C**. PLS-DA of aphids fed on distinct diet diversity levels. R^2^ = 0.61, *P* = 0.185 (ANOVA test). **D**. Classification of aphid chemical features significantly linked to the diversity level of the diet and positively correlated with plant chemistry (*P* < 0.05). *ND*: not determined. Diets included four plants of the same *Tanacetum vulgare* chemotype (C1), two plants of two chemotypes (C2), one plant of four distinct chemotypes (C3), and one plant of three distinct chemotypes plus one plant of the related species *Chrysanthemum indicum* (C4).

## 4. Discussion

Insects evolved mechanisms to neutralise plant defences or even exploit plant metabolites for their own protection against predators or abiotic threats (Beran *et al*. 2018; Singh & Müller 2025). Yet, the extent to which diet shapes the overall insect metabolome as well as chemodiversity remains poorly understood, as studies have focused on specific chemical compounds or families. Is uptake selective for specific compounds, such as amino acids (Broussard *et al*. 2023), or does it broadly reflect plant metabolic diversity? We found that chemical variation in phloem sap-sucking insects not only spans major chemical families but also reflects inter- and intraspecific chemodiversity from plants.

### 4.1. Phloem sap-feeding insects capture plant inter- and intraspecific chemodiversity

Between 78% to 93% of the detected aphid metabolome was shared with their diet, revealing a vast reservoir of potential diet-related chemical influences but also potentially independently shared metabolites. Our results revealed that changing a species or a chemotype in the diet led to profound variation in aphid chemistry. Modifying a species in the diet had consequences on up to 60% of all chemical families used in this study, such as changes in glucosinolate diversity in *M. persicae* and in phenolics and fatty acids in both aphid species. In addition, aphid samples could even be clustered according to the same *T. vulgare* terpenoid chemotype as diet, demonstrating the capacity of phloem sap-sucking insects to capture both inter- and intraspecific variation in their diet. Modifying a terpenoid chemotype influenced phenolics, amino acids, and terpenoids of aphids. These results are consistent with previous studies showing that terpenoid chemotypes of *T. vulgare* vary not only in terpenoids, but also in phenolics and fatty acids (Dussarrat *et al*. 2023). Notably, levels of some chemical features in aphid samples significantly influenced by the diet composition were negatively correlated with the feature level in the diet. The significant variation of these features, combined with the absence of or negative correlation with the feature level in the plant, may arise from metabolic conversion of chemical features within herbivores or de novo synthesis from plant-derived precursors (Friedrichs *et al*. 2022; Hanusch *et al*. 2026). Overall, plant features might be incorporated by the aphids via the phloem sap, which is not only composed of sugars but also transports amino acids, glucosinolates, and fatty acids (Broussard *et al*. 2023; Kos *et al*. 2012; Van Bel 2003). Although less described, some phenolics and terpenoids might also be present in the phloem sap (Bühler & Schweiger 2024; Hijaz *et al*. 2016; Lohaus 2022; Wallis *et al*. 2008). In addition, aphids may capture plant metabolites such as terpenoids or fatty acids via contact with the leaf surface (Batovska *et al*. 2008). For example, the terpenoids of *T. vulgare* are stored in glandular trichomes on the leaf surface (Devrnja *et al*. 2021), which is thus the most likely path for uptake of these metabolites into aphids.

### 4.2. Chemical consequences of a change in diet composition are dependent on time and diet diversity

Our results revealed rapid metabolic turnover in aphids, observable within 24 hours following a change in the plant species composition of their diet. The effect became more pronounced in the following aphid generation(s). This pattern could reflect metabolites that pass through the insect without sequestration (*i*.*e*. metabolites stored and concentrated, Opitz & Müller 2009), leading to greater temporal variability. Detailed studies are needed to test how quickly plant metabolites are turned over and how long diet traces remain in the digestive system of the aphids. Besides temporal variation, chemical consequences of a change in diet composition seemed to be dependent on diet diversity. Chemotypes of *T. vulgare* could be distinguished not only by terpenoids but also by other metabolites. In contrast, adding a chemotype to a diet (*e*.*g*. adding the “Myrox” chemotype from diversity treatment C2 to C3) only led to a partial shift in the overall aphid chemical fingerprint (Fig. 4C). This limited chemical distinctiveness may also be explained by the fact that the same chemotypes (BThu and Aacet) were used in both C1 and C2, and C3 included only two additional chemotypes.

However, this result suggests that the chemical consequences of a loss or gain of a new plant species or chemotype in the aphid diet might be lower on highly diverse diets compared to diets composed of a single or limited number of species or chemotypes. Both time and diet diversity parameters are therefore important to consider when interpreting the results, especially under natural settings. In fact, natural ecosystems offer a broad range of chemotypes (Kleine & Müller 2011), which aphids may mix in their diet, given that wingless individuals (*Aphis gossypii* and *Acrythosiphon pisum*) can walk up to 13.5 meters in 7 hours (Ben-Ari *et al*. 2015; Lombaert *et al*. 2006).

### 4.3. Potential ecological consequences from aphid to ecosystem scale

The ecological consequences of the aphid’s capacity to capture inter- and intraspecific chemodiversity remain to be tested. Herbivorous insects were found to use certain chemicals for their own defence (Müller & Arand 2007). However, since plant chemicals and overall plant quality also influence the preference and performance of herbivores, both phloem-sap sucking aphids and leaf-chewing larvae may face trade-off between benefits (better protection against predators or parasitoids) and costs (lower performance) (Le Guigo *et al*. 2011; Müller & Arand 2007). Studies must now move beyond single metabolites and explore the effect of the overall chemodiversity, as previously described ecological impacts on chemodiversity in plants (Jakobs & Müller 2018; Xiao *et al*. 2025; Ziaja & Müller 2023) may extend to aphids. This hypothesis is supported by previous studies, which demonstrated the importance of plant-derived insect chemical profiles in interactions between insects and parasites or predators (Awater-Salendo *et al*. 2020; Grunseich *et al*. 2021; Müller & Arand 2007).

Results showed that plant inter- and intraspecific chemodiversity escalated to a higher trophic level, raising questions regarding chemical fluxes across the food chain. Are predators also capturing chemodiversity properties of herbivores? How are those chemical movements characterised in plant-soil interactions (Dussarrat *et al*. 2025)? Previous research revealed movements of chemicals across trophic levels, such as pesticides (Wan *et al*. 2025), supporting the possibility of a cascade effect towards predators. Overall, this study highlights the need for a deeper understanding of the extent and ecological consequences of such chemodiversity dynamics across all trophic levels. In addition, results from this study echo the need for chemodiversity conservation, as losing plant chemodiversity might mean losing its functions across the entire trophic chain (Hanusch *et al*. 2026; Le Guigo *et al*. 2011).

## 5. Concluding remarks

This study expanded the herbivore metabolic repertoire potentially influenced by the plant diet. In addition, we found that herbivores can capture plant inter- and intraspecific chemodiversity within short time-scales and across generations. While research on chemodiversity is gaining interest due to its various ecological consequences (Hanusch *et al*. 2026; Petrén *et al*. 2024), our findings represent a significant step forward in our understanding of the dynamics of chemodiversity across the trophic chain. Additional studies are, however, needed to assess the physiological consequences for herbivores and verify whether this concept expands to predators and parasitoids as well as to belowground organisms.

## Acknowledgements

We thank Michael Rostás from Göttingen University for providing us with a starting colony of *M. persicae*. EA, JB and TD thank Mitacs for the Mitacs Globalink Research Award (funding FR143778). JB acknowledges the Natural Science and Engineering Research Council (NSERC, 2019-04516) for financial support. CM and TD are grateful to the German Research Foundation (DFG, project MU1829/28-2) for financial support. We also thank the Open Access Publication Fund of Bielefeld University.

## Conflicts of Interest

The authors declare no conflicts of interest.

## Notes

### Competing Interest Statement

The authors have declared no competing interest.

