## Supplemental Figures for "Aphids capture plant inter- and intraspecific chemodiversity"

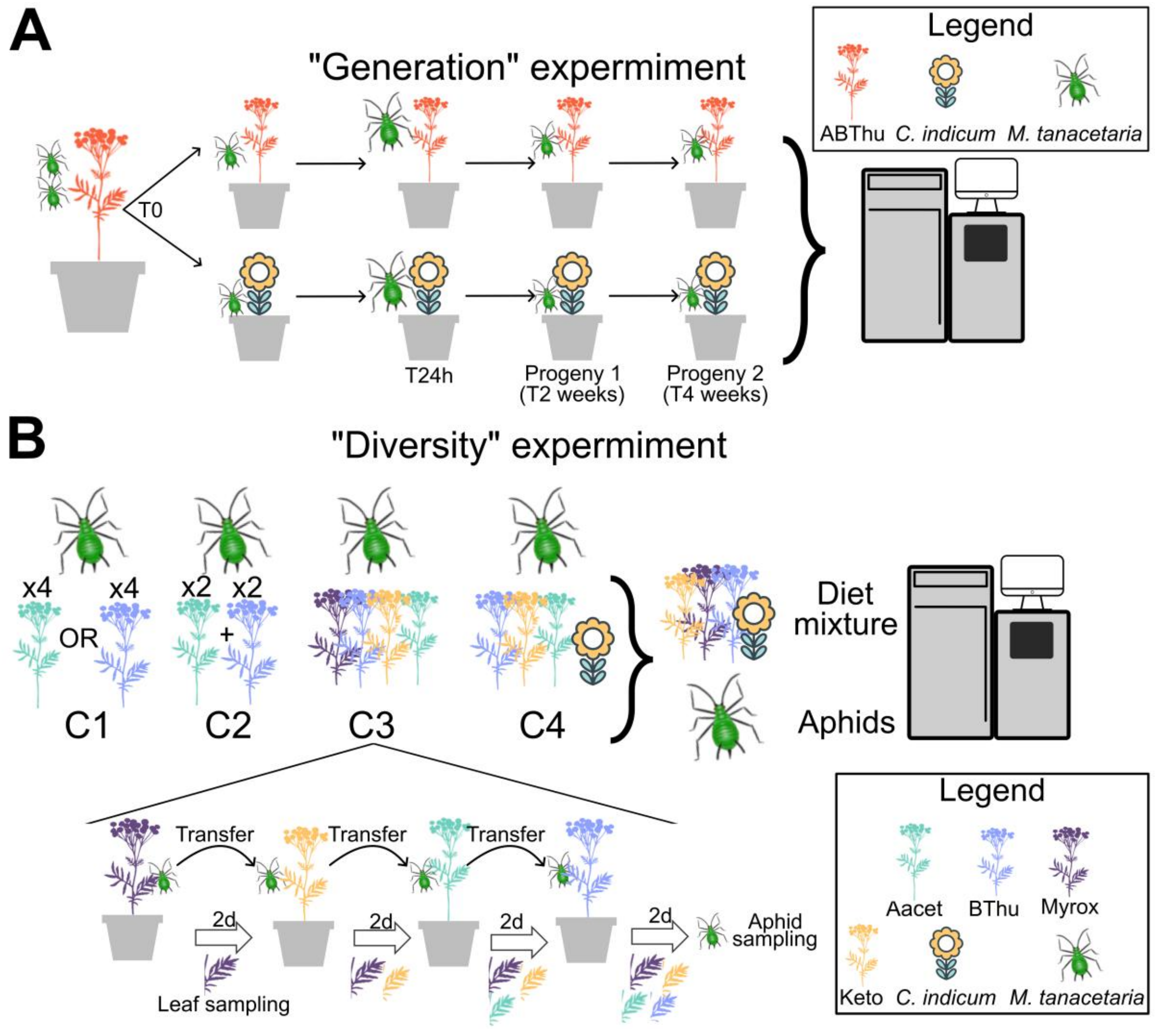


**Fig. S1 | A simplified scheme of the "Generation" and "Diversity" experiments. A.** For the "Generation" experiment, nymphs of *Macrosiphoniella tanacetaria* were placed on potted plants of ABThu (*Tanacetum vulgare* chemotype) at time T0, and 30 aphids were sampled. At time T24 hours, forty to fifty aphids were transferred to *Chrysanthemum indicum* or kept on ABThu, and 30 aphids were collected. Sampling was performed again after two and four weeks. A similar design was performed with *Myzus persicae* on *Brassica oleracea* and *Pisum sativum*. This experiment was done in four replicates. **B.** For the "Diversity" experiment, sixty nymphs were placed into each of four feeding conditions, each comprising four plants on which aphids fed sequentially (C1 to C4). Diets included four plants of the same *Tanacetum vulgare* chemotype (C1), two plants of two chemotypes (C2), one plant of four distinct chemotypes (C3), and one plant of three distinct chemotypes plus one plant of the related species *Chrysanthemum indicum* (C4). Aphids were enclosed in a mesh bag and transferred every two days. Each condition (C1 to C4) included four distinct maternal origins. Both experiments were then processed together into the same LC-MS sequence. This experiment was done with 16 replicates for C1, 16 for C2, 18 for C3, and 14 for C4. *Aacet*: artemisyl acetate-artemisia ketone-artemisia alcohol chemotype, *ABThu*: α-/β- thujone chemotype, *Bthu*: β-thujone chemotype, *Keto*: artemisia ketone chemotype, *Myrox*: (*Z*)-myroxide-santolina triene-artemisyl acetate chemotype. *ND*: not determined.

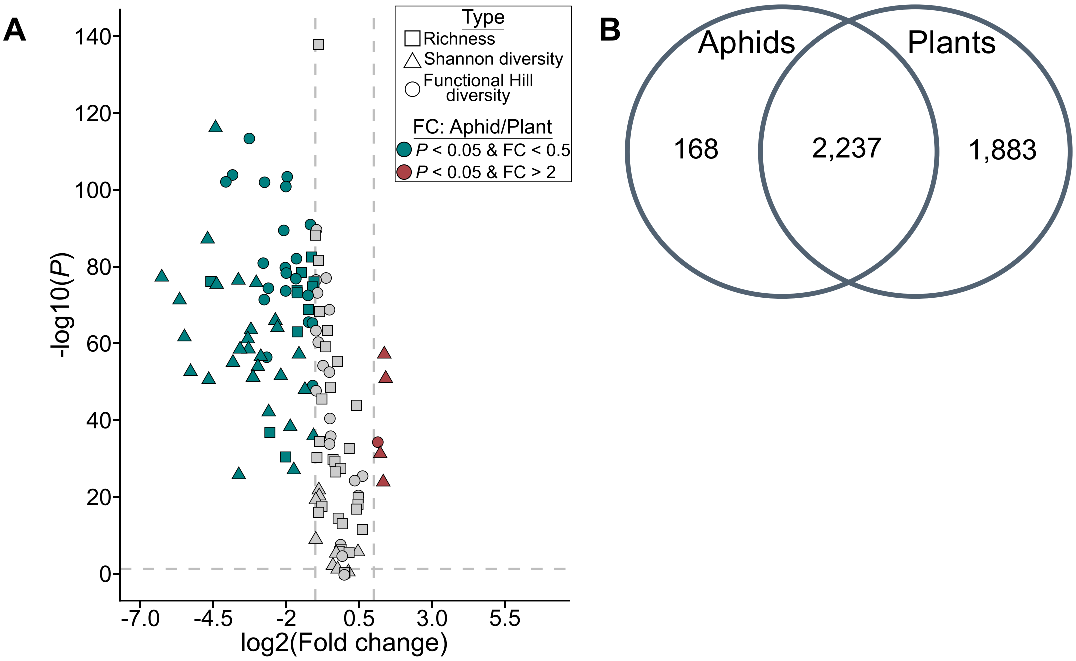


**Fig. S2 | Comparison of aphid and plant samples in the "Diversity" experiment. A.** Depiction of richness, Shannon diversity and Functional Hill diversity in aphids and leaves. Fold changes (FC) were calculated as the ratio between aphids and plants. Blue represented higher values in plants ($P < 0.05$, fold change < 0.5) while red represented higher values in aphids. **B.** Detected metabolic features in aphids and plants.

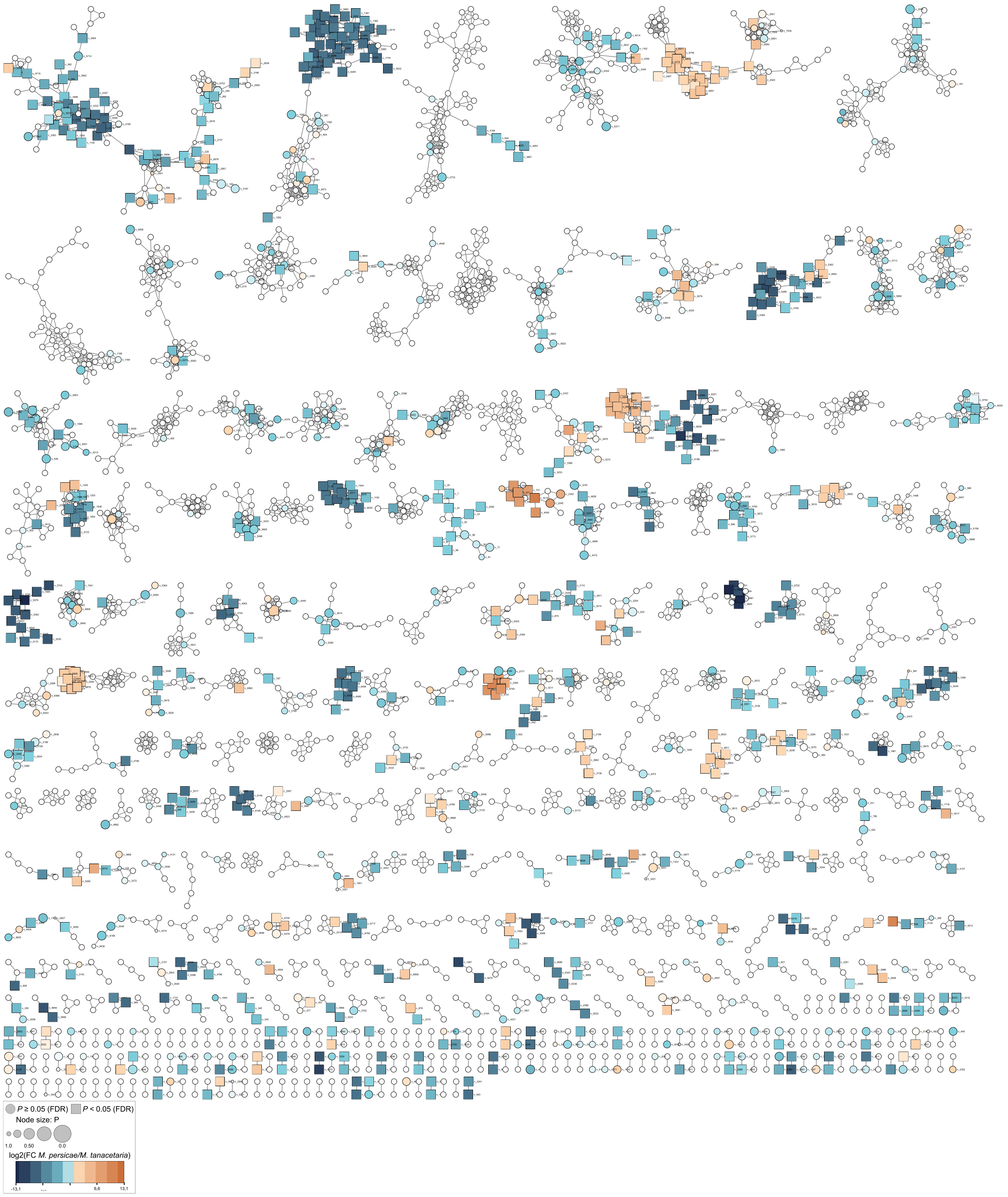


**Fig. S3 | Feature-Based Molecular Network depicting major chemical variation between *M. persicae* and *M. tanacetaria*.** Brown colour refers to higher contents in *Myzus persicae*, while blue colour refers to higher contents in *Macrosiphoniella tanacetaria*. *FC*: fold change.

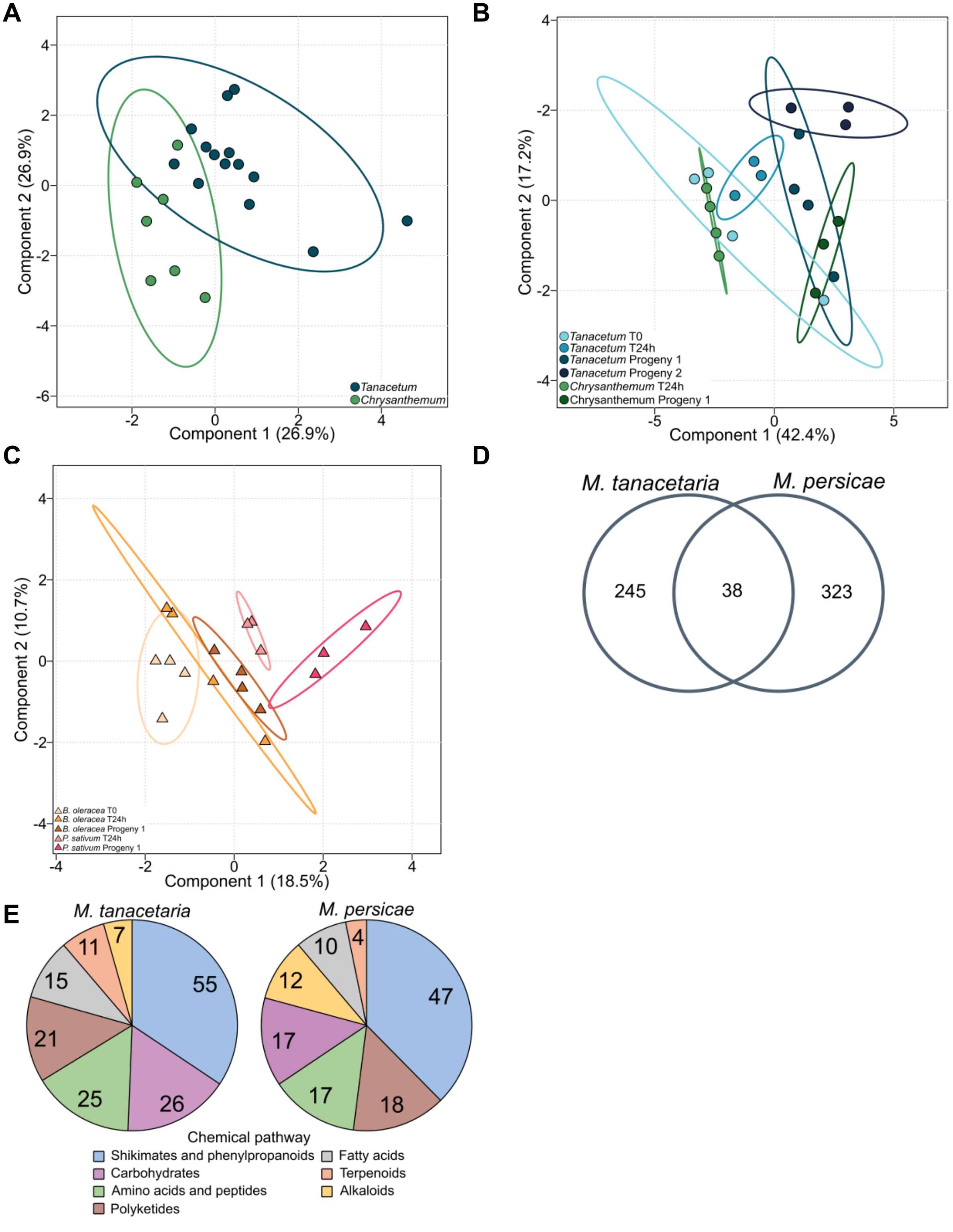


**Fig. S4 | Aphid chemical features influenced by the plant diet in *M. tanacetaria* or *M. persicae*. A**. PLS-DA exploring the effect of diet on chemodiversity indices of *Macrosiphoniella tanacetaria* without considering the generation effect. $R^2 = 0.66$, $P = 0.387$ via ANOVA test. **B.** PLS-DA exploring the effect of diet on chemodiversity indices of *Macrosiphoniella tanacetaria* through asexual generations. $R^2 = 0.66$, $P_{species} = 0.169$, $P_{generation} < 0.001$, $P_{species*generation} = 0.257$ via ANOVA test. **C.** PLS-DA exploring the effect of diet on chemodiversity indices of *Myzus persicae* through asexual generations. $R^2 = 0.90$, $P_{species} = 0.004$, $P_{generation} = 0.007$, $P_{species*generation} = 0.610$ via ANOVA test. **D.** Depiction of chemical features significantly influenced by the plant species included in the diet ($P < 0.05$) in *M. tanacetaria* and *M. persicae*. **E.** Putative annotation of significant chemical features.

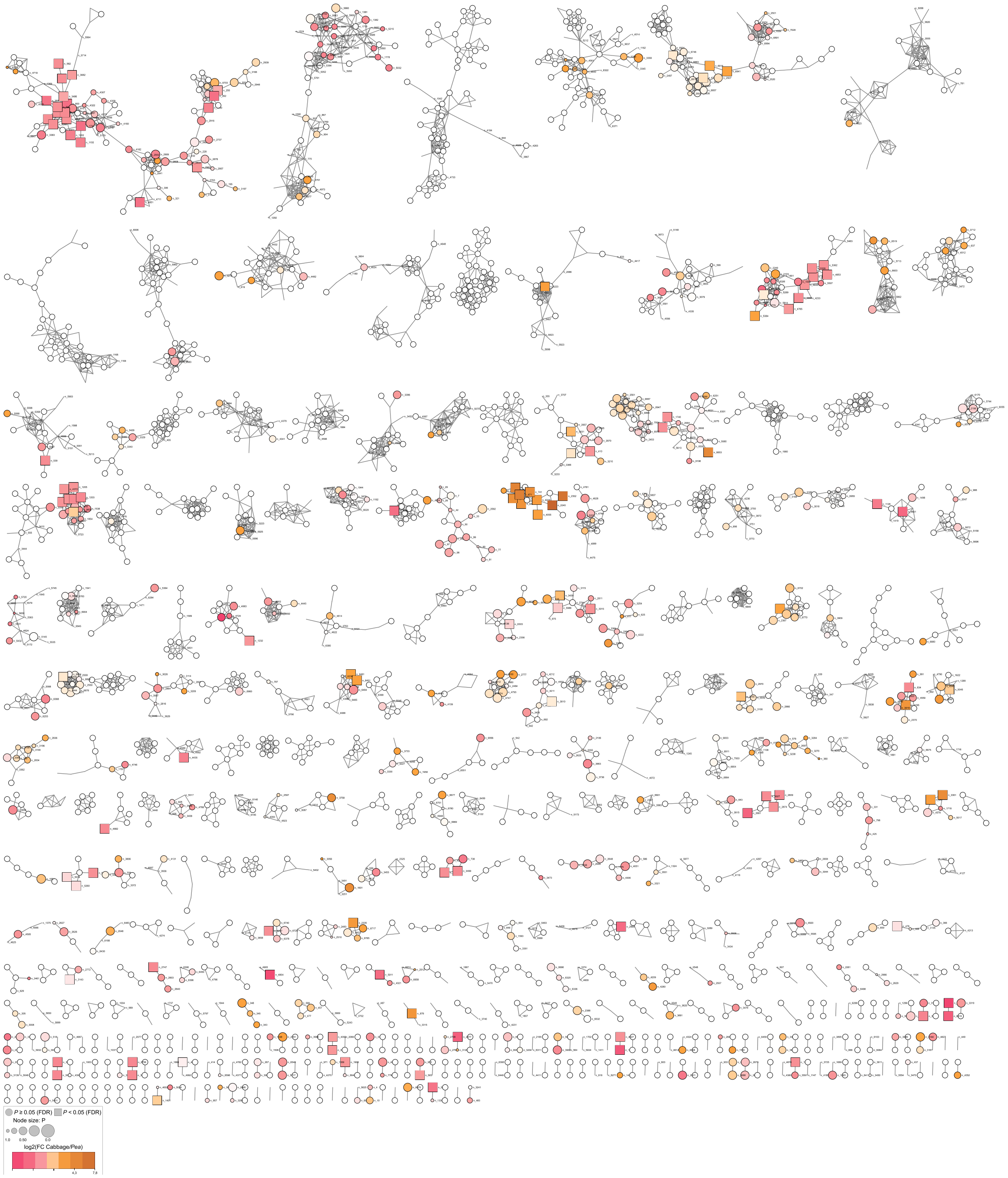


**Fig. S5 | Feature-Based Molecular Network depicting major chemical variation between *M. persicae* feeding on *B. oleracea* or *P. sativum*.** Brown colour refers to higher contents in aphids feeding on *B. oleracea* (cabbage), while pink colour refers to higher contents in aphids feeding on *P. sativum* (pea). *FC*: fold change (*B. oleracea*/*P. sativum*).

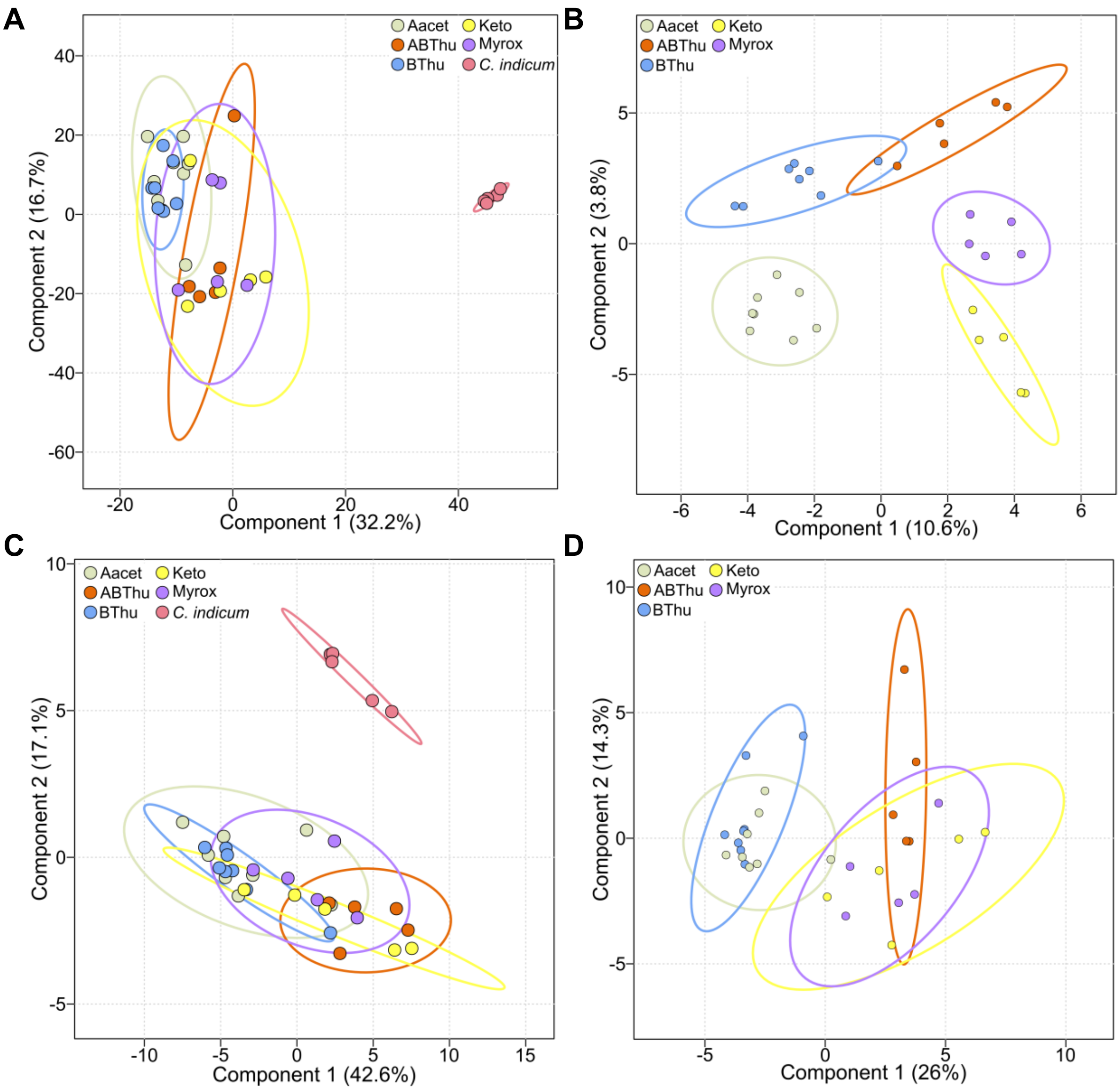


**Fig. S6 | Dilution effect of diverse diet. A-B.** Principal component analysis (PCA, **A**) and sparse partial least squares-discriminant analysis (sPLS-DA, **B**) of leaf metabolism (4,288 features) comparing chemotypes. Classification error rate of the sPLS-DA is equal to 12.9% (5-fold CV). **C-D.** PCA (**C**) and sPLS-DA (**D**) of leaf chemical indices (126). Classification error rate of the sPLS-DA is equal to 64.5% (5-fold CV). *Aacet*: artemisyl acetate-artemisia ketone-artemisia alcohol chemotype, *ABThu*: α-/β- thujone chemotype, *Bthu*: β-thujone chemotype, *Keto*: artemisia ketone chemotype, *Myrox*: (*Z*)-myroxide-santolina triene-artemisyl acetate chemotype. sPLS-DA were built with 20 features per component. *Chrysanthemum indicum* was removed from the sPLS-DA to explore the variation between chemotypes.

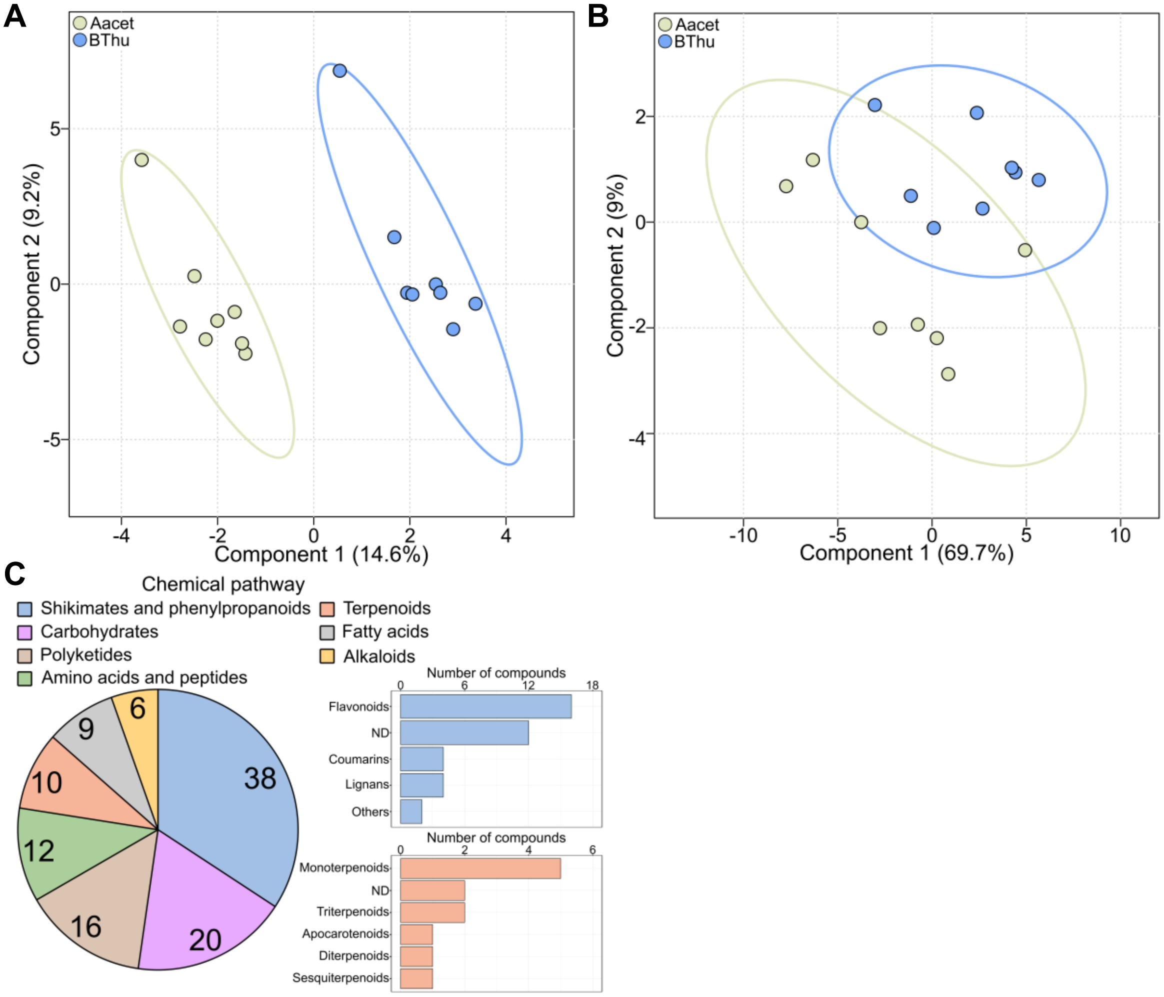


**Fig. S7 | Discriminant capacity of aphid metabolome. A.** Sparse partial least squares-discriminant analysis (sPLS-DA) comparing the metabolome of aphids that fed on β-thujone chemotype (BThu) or artemisyl acetate-artemisia ketone-artemisia alcohol chemotype (Aacet*)*. sPLS-DA models were defined using 20 features per component. Classification error rate of the sPLS-DA is equal to 18.8% (5-fold CV). **B.** Partial least squares-discriminant analysis (PLS-DA) comparing aphid chemical indices (126 indices) that fed on BThu or Aacet chemotype. $R^2 = 0.67$, $P = 0.122$ (ANOVA test). **C**. Classification of aphid chemical features significantly linked to the diversity level of the diet ($P < 0.05$). *ND*: not determined.
